# A global envelope test to detect early and late bursts of trait evolution

**DOI:** 10.1101/175968

**Authors:** D. J. Murrell

## Abstract

The joint analysis of species’ evolutionary relatedness and their morphological evolution has offered much promise in understanding the processes that underpin the generation of biological diversity. Disparity through time (DTT) is a popular method that estimates the relative trait disparity within and between subclades at each time point, and compares this to the null hypothesis that trait values follow an uncorrelated random walk along the time calibrated phylogenetic tree. A simulation envelope is normally created by calculating, at every time point, the 95% minimum and 95% maximum disparity values from multiple simulations of the null model on the phylogenetic tree. The null hypothesis is rejected whenever the empirical DTT curve falls outside of this envelope, and these time periods may then be linked to events that may have sparked non-random trait evolution. However, this method of envelope construction leads to multiple testing and a poor, uncontrolled, false positive rate. As a consequence it cannot be recommended. A recently developed method in spatial statistics is introduced that constructs a confidence envelope by giving each DTT curve a single ranking value based upon its most extreme disparity value. This method avoids the pitfalls of multiple testing whilst retaining a visual interpretation. Results using simulated data show this new test has desirable type 1 properties and is at least as powerful in correctly rejecting the null hypothesis as the morphological disparity index and node height test that lack a visual interpretation. Three example datasets are reanalyzed to show how the new test may lead to different inferences being drawn. Overall the results suggest the new rank envelope test should be used in null model testing for DTT analyses, and that there is no need to combine the envelope test with other tests such as has been done previously. Moreover, the rank envelope method can easily be adopted into recently developed posterior predictive simulation methods. More generally, the rank envelope test should be adopted when-ever a null model produces a vector of correlated values and the user wants to determine where the empirical data is different to the null model.

## Introduction

Understanding the joint temporal dynamics of taxonomic and phenotypic diversity can provide tremendous insights into evolutionary success and its relationship with ecological opportunity, selective pressures, constraints, biotic interactions and environmental conditions. At the most basic level evolutionary biologists are often interested in detecting non-random evolution of biological traits within and across clades of species. Non-random bursts in evolution are often thought to be associated with events that open up ecological opportunities and enable a rapid increase in speciation rates and trait evolution, followed by slowdown in both processes as the ecological niches become filled. The evolutionary theory of adaptive radiation is the special case where the burst in speciation rate and trait evolution occur early in the clade’s history (Schluter, 2000), but such bursts in trait evolution may occur at other times and can be triggered by other processes such as major events in the external environment.

A variety of methods exist to look for the signature of evolutionary bursts and fall into the categories of null model testing, and model selection (Harmon et al., 2003, Freckleton and Harvey, 2006, Harmon et al., 2010, Slater et al., 2010, Slater and Pennell, 2014). The model selection approach takes a variety of candidate models (Brownian evolution, early burst, selective peak) and fits these to the data using maximum-likelihood methods before choosing the model that has the ‘best fit’ (Harmon et al., 2010, Slater and Pennell, 2014). The null model approach remains more popular, partly because the methods have been established for longer, and the overall aim is to investigate if the data can be distinguished from the null model of uncorrelated evolution of trait values (Harmon et al., 2003, Freckleton and Harvey, 2006).

One of the more popular null model approaches is to look at morphological traits to see if trait disparity increases, decreases or stays the same as species accumulate in evolutionary time, and also see whether this disparity is greater within clades or between clades. Convergent evolution of traits is implied if morphological disparity is predominantly found within one or more subclades; whereas adaptive radiations are expected to show divergence of traits between subclades, and in this scenario between clade morphological disparity should be greater than among subclade disparity. This analysis of between and within clade trait disparity has been championed by the disparity through time (DTT) approach introduced by Harmon et al. (2003). Here the empirical DTT curve is compared to the distribution of DTT curves generated on the same phylogenetic tree but under a specific model of how the trait diversity evolves. Generally the null model is an uncorrelated random walk, and this is generally referred to as Brownian evolution (ie a Brownian random walk over time in trait space). The method of comparison is critical in determining whether the empirical data can be distinguished from the null model. Early analyses used an integral deviation method called the Morphological Disparity Index (MDI) which sums the deviations of the empirical DTT curve from the median of the null model simulations (Harmon et al., 2003). The index can then be compared to the distribution of values produced by the simulation to test whether it is significantly different from the null model (e.g. Ingram (2015)). Where MDI > 0, this implies within-clade trait variation is generally greater than expected under the null model, and MDI < 0 implies between-clade trait variation is more dominant than expected under the null model, and is suggestive of an adaptive radiation. The strength of the MDI is that it is a global test and so avoids multiple testing that can occur in analyses of time series data (see below), but the power of the MDI to detect non-Brownian bursts in trait evolution may be compromised if short-lived but extreme deviations in one direction at one point in the time series are balanced by a weak but long-lived deviation in the opposite direction.

Since the MDI produces a number, visualization of where any non-random bursts might have occurred (e.g. early on in the radiation) can only proceed by plotting the empirical DTT curve against the DTT curves sampled from the null model. However, determining where *statistically significant* local deviations from the null model are occurring in the time series requires another test. Slater et al. (2010) provided this by introducing an envelope method where the (100-α)% upper and lower confidence intervals for the null model are estimated by sampling from the null model *n* times (where typically, *n* > 1000) and then ordering the relative disparity values at each time point to arrive at an envelope that contains (100-2α)% of the simulated relative disparity values at each time point. This method is also referred to as the pointwise envelope method (Myllymaki et al., 2017). The observed relative disparity can then be compared to this envelope and if it falls outside the null model is said to be rejected at that level of significance.

The pointwise envelope method continues to be a popular method of inference. For example, Weber et al. (2016) used the DTT method to investigate the macroevolution of perfume signalling in orchid bees, a group known for their chemical sexual communications, and therefore a likely candidate for rapid diversification in traits. They found strong non-Brownian evolution for perfume signal, and weaker support for non-Brownian evolution of the labial gland. In both cases disparity was greater than the median DTT of the null model simulations indicating clades are overlapping in trait space, a signature of convergent evolution. The visual/graphical interpretation of the DTT with an envelope test has extra appeal as it can be used to identify time points where the burst of non-Brownian evolution occurred, enabling correlation with known evolutionary or environmental events that have triggered the burst. For example, Aristide et al. (2016) investigated brain shape, encephalization, and log body mass in New World monkeys. Their DTT envelope analyses supported the conclusion that a burst in evolution of brain shape occurred approximately 17-12 Ma, was associated with a burst in evolution of body mass that has previously been linked to diversification of diet and locomotion strategies, and was followed by a slowdown in disparity changes that persists to the current day. Conversely, Feilich (2016) found disparity in cichlid fin and body morphology was often greater than expected under the null model of Brownian evolution indicating most variation in morphology occurred within subclades. Moreover, the observed body and median fin disparity above the 95% confidence interval produced by the null model simulations coincided with the Cichlinae–Pseudocrenilabrinae split, and a later split caused by the radiation of the haplochromine cichlids.

However, the pointwise envelope method leads to weaker than expected statistical performance because multiple tests, one at each time point, are being performed simultaneously. This is an issue that occurs in many different areas (eg spatial statistics Baddeley et al. (2014)), and generally whenever the pointwise envelope method is used in conjunction with a non-parametric method that produces a function as its summary output. Multiple testing leads to an increased type 1 statistical error rate (an elevated rate of rejection of the null hypothesis when it is true) that is no longer in line with the significance level being used to generate the confidence intervals of the envelope. Although multiple testing problems may be solved using a Bonferroni correction, it is not appropriate here because the assumption of independence of tests is violated by the correlation of disparity values between consecutive time points and also the (often) large number of time points being simultaneously evaluated (Loosmore and Ford, 2006). Perhaps as a consequence of this many studies, including those discussed above, have used multiple methods to look for non-Brownian trait evolution including the MDI and the node height test (Freckleton and Harvey, 2006). However, the continued use of the pointwise envelope suggests its graphical interpretation is very appealing and it would therefore be worthwhile to circumvent its multiple testing issues.

Recently, a new method that avoids the multiple testing problems of the pointwise envelope but retains the visual interpretation has been developed in spatial statistics (Myllymaki et al., 2017). Spatial analysis of ecological data often leads to use of a non-parametric summary statistic such as Ripley’s K that plots the tendency to cluster against the radial distance (eg. Law et al. (2009)), and the problem of pointwise envelopes for inference of non-random patterns is well established (Loosmore and Ford, 2006, Baddeley et al., 2014). Instead of producing confidence intervals by ranking each curve at each time point the rank envelope method ranks each curve for its overall extremness (more details are given below). As shown by Myllymaki et al. (2017), it has good type 1 and type 2 error rates and is recommended for testing point pattern data against the null model of complete spatial randomness. The rank envelope can be developed and applied to any model that produces a vector (eg van Veen and Murrell, 2005), however, its performance needs to be tested since there are many ways of ordering curves based on their ‘extremness’ and not all methods will produce desirable results.

In what follows, the rank envelope test will be developed for DTT null model analyses and its type 1 and type 2 statistical properties compared to the pointwise envelope, MDI, and node height tests. The pointwise envelope test will be shown to have extremely poor type 1 error rates and should not be used for inference. In contrast, the rank envelope method will be shown to possess desirable type 1 error rates, and be at least as powerful as the other tests in detecting accelerating or decelerating rates of trait evolution whilst retaining the useful property of graphical interpretation.

## Methods

### Data simulation

Phylogenetic trees were generated within *R* (version 3.3.3) using the *pbtree* function with the *phytools* R library (version 0.6, Revell (2012)), using the pure-birth (Yule) model. A pure-birth model is a simple way to ensure the number of tips is constant across simulations, but may be biologically plausible for clades over relatively short periods of time. These phylogenetic trees were then used to simulate quantitative trait evolution under a variety of scenarios including the null model of Brownian evolution. Specifically, trait evolution was simulated using the *fastBM* (in *phytools*) and *rescale* (in *geiger*, version 2.0.6, Pennell et al. (2014)) functions. The *rescale* function allows the simulation of early or late burst trait evolution using the early burst option, and a rate change parameter, *a*. When *a* < 0 the rate of evolution decreases with time, mimicking an early burst of trait evolution, whereas *a* > 0 models a late burst. The magnitude of *a* determines how quickly this burst of activity fades away or builds up, with large magnitudes delivering a rapid decay or late increase in evolutionary change (examples are given in Figure S1 of the supplementary information). The null model of Brownian evolution is simulated under the same framework, but where *a* = 0.

### Disparity Through Time (DTT) Analyses

DTT has proven to be one of the more popular approaches and uses the average pairwise Euclidean distance between species trait values as a measure of disparity. Following Harmon et al. (2003) relative disparity is calculated by dividing disparity of each subclade by the disparity of the whole tree. At each time point (speciation event) the average relative disparity for that time point is calculated as the mean of the relative disparities for all subclades whose ancestral lineages are present at that time. Disparity values close to zero indicate that variation in the trait(s) is predominantly partitioned between subclades rather than within them. Disparity values close to unity suggest that a clade contains a large amount of that variation, and that clades may overlap in trait space. By definition, disparity is 1 at the base of the phylogenetic tree, but is 0 at the present day.

Two methods that are currently used to search for the signal of bursts in morphological evolution using DTT are (1) the pointwise envelope test (Slater et al., 2010), and (2) an integral deviation test known as the Morphological Disparity Index (MDI, Harmon et al. (2003)). As well as these a third test, the global envelope test, was investigated. The global envelope test retains the same visual interpretation as the pointwise envelope test whilst avoiding the problem of multiple testing. All make comparisons of the empirical DTT to the DTT taken from the ensemble of simulations generated by the null model of Brownian evolution, and all use the same measure of disparity defined above.

### The Pointwise Envelope Test

The pointwise envelope test is a Monte-Carlo simulation method that aims to produce a confidence interval, or envelope within which any part of the empirical DTT curve is said to be statistically indistinguishable from the null model. The method currently implemented in *Geiger* (v2.0.6) constructs an envelope by extracting the αth and (100-α)th quantile of the DTT at each time point (speciation event) for all the null model simulations (normally α = 2.5). This produces a lower and upper interval within which (100-2α)% of all values for DTT at each speciation event under the null model. More formally, the envelope is defined by the lower and upper bounding curves

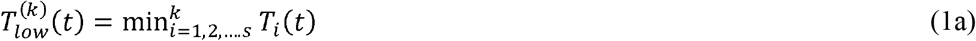

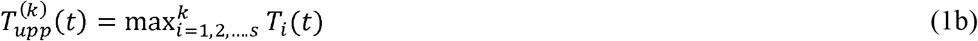

where min^*k*^ and max^*k*^ denotes the *k*th smallest and largest values of the DTT across all simulations *s,* of the null model at time (speciation event) *t*. If the empirical DTT curve falls outside of this envelope it is interpreted as being evidence for a departure from the null model of Brownian evolution.

### The MDI test

Perhaps the simplest way to avoid multiple testing is to perform a deviation test that sums the deviations of the empirical DTT from the median DTT of the ensemble of null model simulations. Known as the Morphological Disparity Index (MDI), negative values indicate the empirical DTT curve is below the null model median DTT for at least some of the range of time points, again pointing to the possibility of an early burst in diversity (Harmon et al., 2003). A disadvantage of this approach is that it is not possible to say where the empirical DTT deviates from the null model without plotting it against the null model simulations and then performing some sort of envelope test. Moreover, since the index sums up the deviations from the median of the ensemble of simulations of the null model, it is theoretically possible for time periods where the empirical DTT is above the median DTT to be cancelled out by time periods where it is below the median DTT, thus giving an MDI value close to that expected under Brownian evolution. However, most likely because it avoids the issue of multiple testing, the MDI has been well used (eg Harmon et al. (2003), Slater et al. (2010), Colombo et al. (2015), Ingram (2015), Jonsson et al. (2015)). In the results below Monte Carlo simulations were used to generate a distribution of MDI values from the null model. The empirical MDI value is then compared to this distribution and the null hypothesis (that the empirical data is from the same distribution as the null model of Brownian evolution) is rejected if it is smaller than the αth quantile value or larger than the (100-α)th quantile. As for the envelope tests, α = 2.5.

### Rank Envelope Test

A method that avoids multiple testing and therefore one which should show better type 1 error properties is the rank envelope test. In this case the whole DTT curve is ranked in terms of its extremness (rather than rank each speciation event individually). This so-called ‘extreme rank’ depth measure method was recently introduced by Myllymaki et al. (2017) for a similar problem that occurs in hypothesis testing for spatial point processes where non-parametric functions (pair correlation function and Ripley’s K) are used to detect departures from complete spatial randomness (ie a lack of any correlation between points). Such tests have been popular in ecology (e.g. Wiegand and Moloney (2004), Flügge et al. (2012)). The more formal underpinnings of the test can be found in Myllymaki et al. (2017), but briefly the DTT curves from the *r* simulations of the null model are ordered according to the largest *k* for which they are still present in the *k*th envelope as defined by the pointwise envelope (equations 1a and 1b).

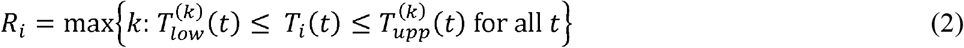

*R_i_* is therefore the most extreme of the *t*-wise ranks of the curve *T_i_(t)*. In other words, each curve is ranked relative to the other simulation curves, with its ranking being taken as its most extreme ranking within the pointwise envelope.

Since ties are possible (indeed are likely) the set of curves can only be weakly ordered. In order to generate a *p*-value a method for dealing with the ties needs to be used. Following Myllymaki et al. (2017), a range of *p*-values is reported that encompasses the most liberal and most conservative *p*-values respectively defined as

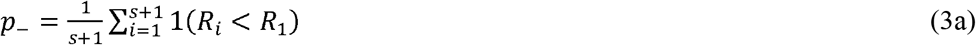

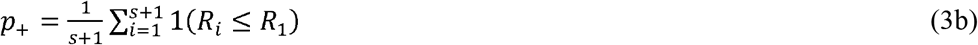

and where *R_1_* is the rank of the empirical DTT curve. This does raise the further problem that the interval defined by *p_-_* and *p_+_* could include the significance level α, leading to an ambiguous result. However the likelihood of this happening is very small as long as *s*, the number of Monte Carlo simulations of the null model is sufficiently large. Myllymaki et al. (2017) recommend *s* ≥ 2500, and the results below use *s* = 5000. From the above DTT curve ordering, it is straightforward to visualise the global envelope determined by the significance level used, and in so-doing the user can readily see where the empirical data falls outside of the global envelope, and thus where the DTT is significantly different to Brownian evolution.

The extreme depth ranking method used here is but one of a number of possible ways to order the DTT curves. Some other functional depth orderings are discussed in Myllymaki et al. (2017) and the reader is directed to their paper for more details, but as will be shown below, the extreme rank depth measure described above leads to desirable type 1 and type 2 statistical errors for DTT analyses.

### The Node Height Test

The final test does not use simulations of the null model to compare to the empirical data but instead relies upon the expectation that trait evolution should slow as niche space become packed. The node height test (Freckleton and Harvey, 2006) investigates if there is a significant correlation between the absolute magnitude of the standardized independent contrasts of the trait(s) and the height above the root of the node at which they were being compared to. The height of a node is defined as the absolute distance between the root and the most recent common ancestor of the pair from which the contrast is generated. A significant relationship between these indicates that the rate of trait evolution is changing systematically through the tree with early and late bursts in trait evolution being diagnosed by the sign of the slope. Since this is an established test, the analyses of the node height test were performed using the function *nh.test* with the R library *geiger* (version 2.0.6).

### Software code

All results were obtained using *R* (version) and the code used to produce the results reported below can be found at https://github.com/djmurrell/DTT-Envelope-code. Details of the current methods to detect non-Brownian bursts of trait evolution can be found in the *geiger* software library, and the rank envelopes are produced by modifying code from the *spptest* library (Myllymaki et al., 2017).

## Results

### False positive rates (Type 1 errors)

The false positive rate is investigated by simulating an empirical dataset of trait evolution under the Brownian null model and testing how frequently each of the four tests described above incorrectly rejects the null hypothesis. Results for the DTT approach using the pointwise envelope test at the 5% level of significance show a disappointing, but unsurprising high rate of false positives (Figure 1). The multiple testing nature of this method means that the false positive rate is dependent upon the number of species in the comparison, and in the simulations the rate of false positives ranges approximately between 0.25 and 0.5 for 10-200 species (Figure 1). That is to say, for comparisons using more than 100 species the pointwise envelope test is incorrectly rejecting the null hypothesis of Brownian evolution in approximately 50% of cases. As such it is impossible to recommend this method for inference of non-Brownian bursts of trait evolution. In comparison, the node height test and the global rank envelope test both return consistent false positive rates that hover around the significance level used (Figure 1). The MDI shows a high type 1 error rate for phylogenetic trees with less than 40 species, but thereafter a desirable false positive rate is returned.

**Figure 1.**
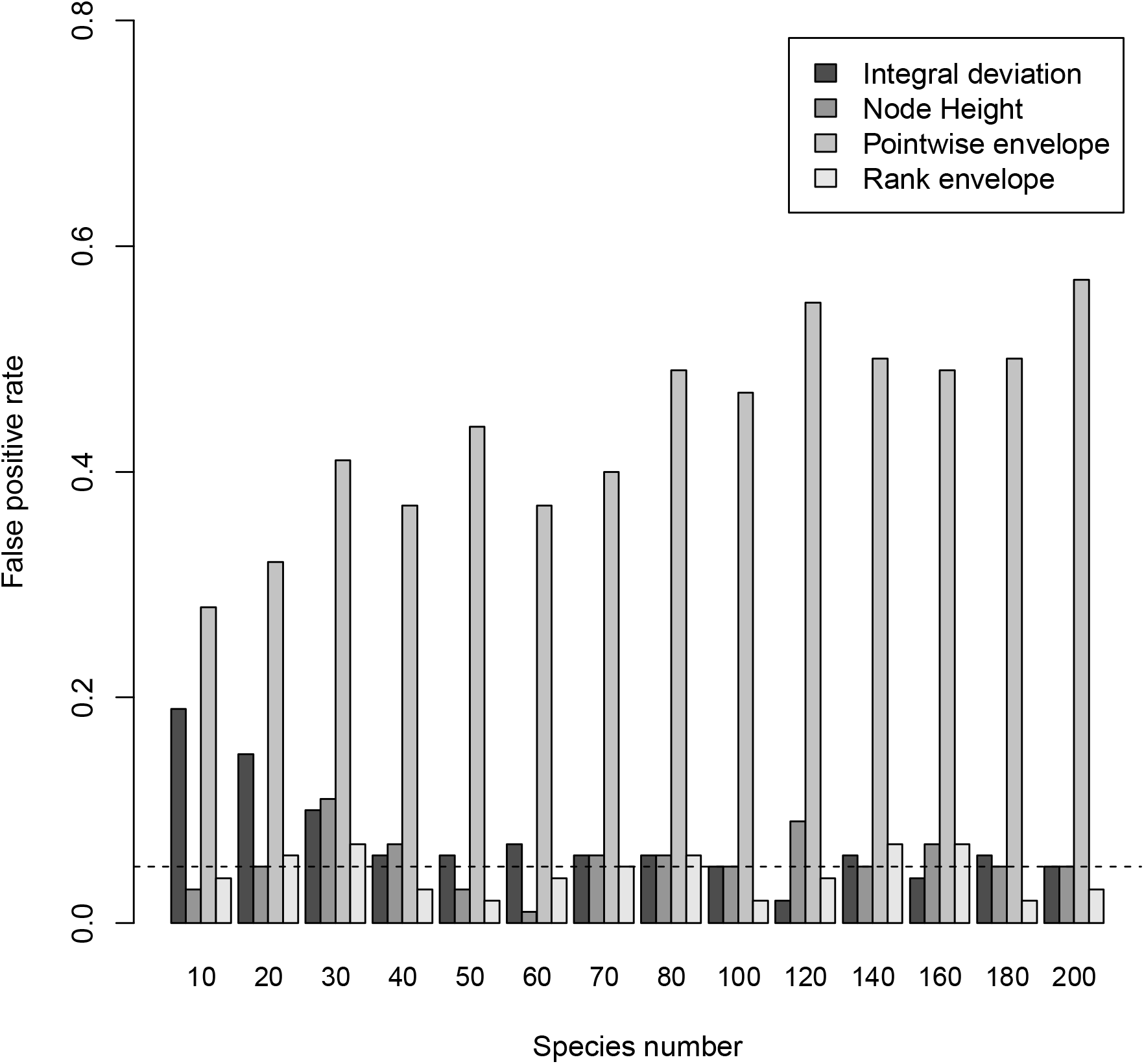
False positive rates for the four tests for non-random disparity through time (DTT) as a function of the number of species at the tips of the phylogenetic tree. Rates are estimated from 100 simulated phylogenetic trees for each number of species using a pure birth model to generate the phylogenetic tree, and assuming Brownian evolution of the trait at each speciation event. All tests requiring Monte Carlo simulations were run with *s* = 5000 trait evolution simulations.

### True positive rates (Type 2 errors)

Evolutionary biologists are often most interested in early bursts in trait evolution, since this is argued to be the hallmark of adaptive radiations (Harmon et al., 2010). However, a number of studies have also reported late bursts in trait evolution where departures from Brownian evolution occur only in the recent past (e.g. Koecke et al. (2013), Tran (2014), Pincheira-Donoso et al. (2015), Feilich (2016)). Simulations for both scenarios confirm that the MDI, the node height, and the rank envelope test can all successfully detect both early and late bursts in trait evolution (Figure 2). Convention dictates that at the 5% significance level a desirable test shows a true positive rate of 0.8. The ability of all tests to reach this mark is dependent on the number of species and the strength of the early or late burst, but other generalities do emerge. Firstly, for early burst trait evolution the MDI deviation test is less powerful than the node height and global envelope tests, with the global envelope test generally showing slightly higher power. Secondly, early bursts are slightly easier to detect than late bursts of trait evolution i.e. the desired true positive rate is reached with fewer species in early burst models compared to late burst models. The node height and global envelope tests show similar power to detect late bursts, but the MDI is better able to detect late bursts in trait evolution with fewer species at the tips. Example simulations for each early/late burst rate with 100 species at the tips of the phylogenetic tree can be found in the supplementary material (Figure S1).

**Figure 2.**
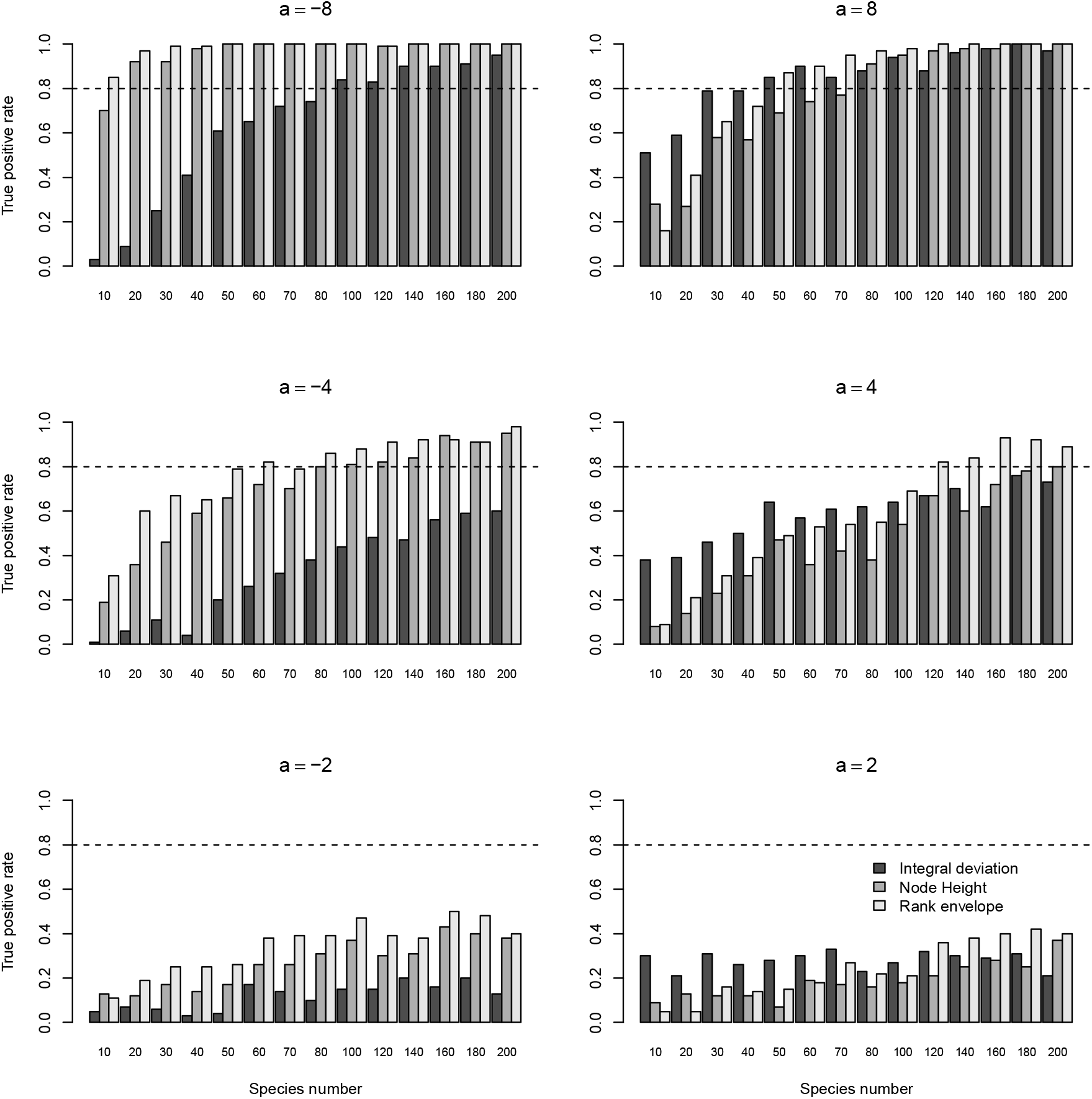
True positive rates (statistical power) of the rank envelope test, the MDI test and the node height test under a range of simulated early and late burst scenarios, and for a range of size of phylogenetic tree. Rates are estimated from 100 simulated phylogenetic trees for each number of species using a pure birth model to generate the phylogenetic tree, and assuming trait evolution at each speciation event speeds ups or slow downs over evolutionary time. All tests requiring Monte Carlo simulations were run with *s* = 5000 trait evolution simulations for every phylogenetic tree. When *a* < 0, a burst in trait evolution occurs early in evolutionary time, and late bursts occur when *a* > 0. Large magnitudes lead to the bursts occurring over a smaller period of time. Corresponding example plots of DTT in each scenario are given in the supplementary information.

### Data Examples

Having established the rank envelope test possesses desirable type 1 and type 2 statistical error properties, three datasets were used to illustrate how inference of the rates of morphological evolution can change depending on whether the pointwise or global envelope test is used. Since the pointwise envelope test is too liberal in its rejection of the null model the expectation should be for a reduction in support for non-Brownian bursts in morphological evolution.

The first example uses the morphological and phylogenetic data on Darwin’s finches (Geospiza) which is currently found in the *geiger* (version 2.0.6) *R* package. Re-analysis shows support for two late bursts in culmen length evolution by the pointwise envelope test, both showing diversification predominantly occurring within clade(s) followed by decreases in disparity caused by increased diversity between clades (Figure 3a). In contrast, there is no departure from the null model of Brownian evolution according to the rank envelope test (Figure 3b).

**Figure 3.**
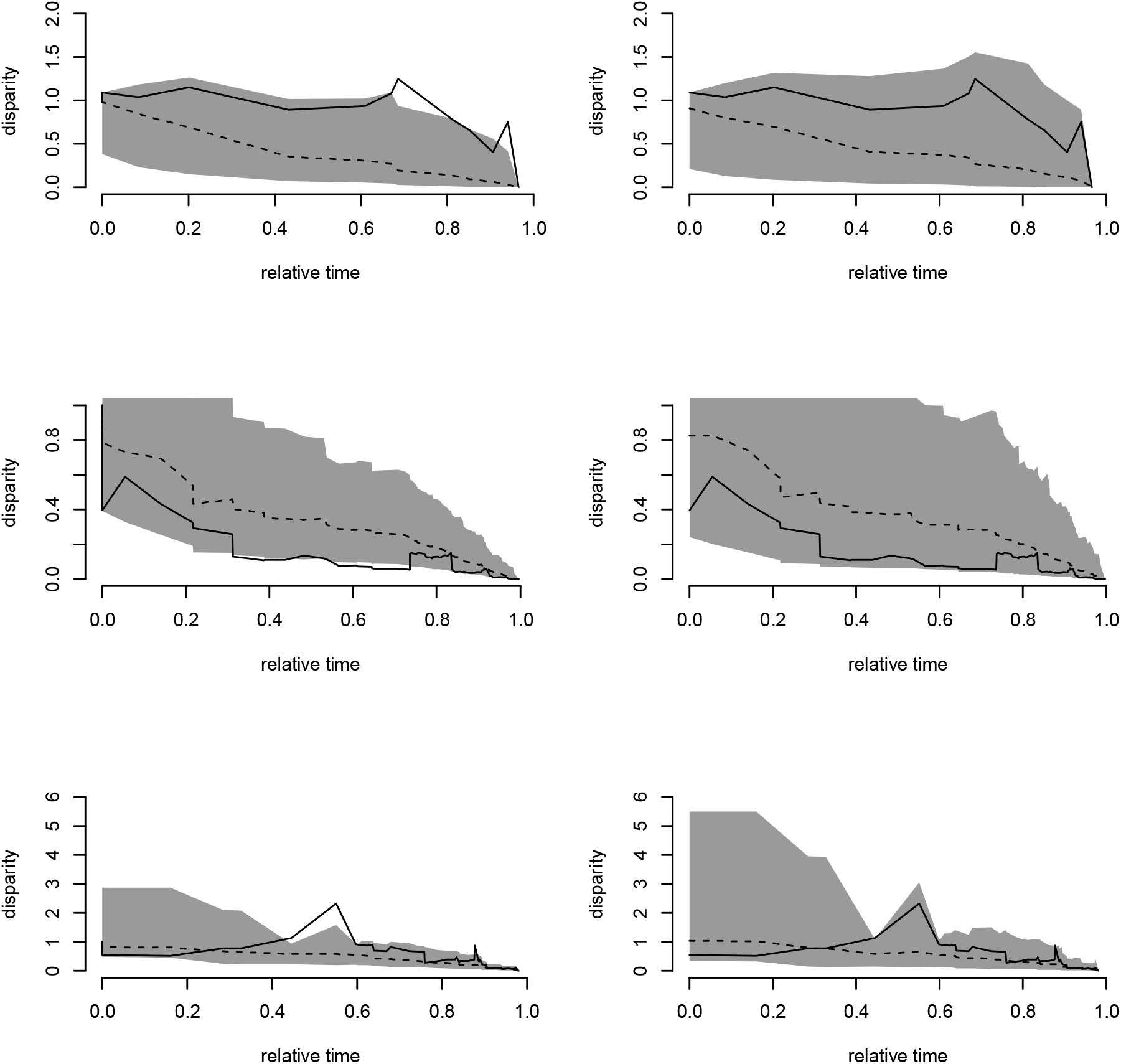
Comparisons of inference from using the pointwise envelope test (left hand column) and the rank envelope test (right hand column) for trait disparity through time for three showcase datasets. In each panel, the empirical pattern (solid black line) is compared to the median of 5000 simulations of the null model of Brownian evolution (broken lines), and the shaded regions correspond to the 95% confidence intervals calculated using the pointwise (left column) and rank envelope (right column) methods. Top row is for Darwin’s finches (Geospiza) and the evolution of culmen length; middle row is for the evolution of Cetacean body size (Slater et al., 2010); bottom row is for the evolution of anal fin shape in 131 species of African cichlids (Feilich, 2016).

The second example uses a time-calibrated molecular phylogeny of extant cetaceans and a morphological dataset on body size from Slater et al. (2010) which is also located within *geiger* (version 2.0.6). Their analyses used a combination of the node height test (NHT), and DTT tests using the MDI and the DTT pointwise envelope method. On the basis of the pointwise envelope results they came to the conclusion that cetaceans do show a burst in evolution of body size and that this occurred predominantly during the period 6-11 Ma. Reanalysis confirms that the pointwise envelope approach finds the same burst in trait evolution but that the global envelope test fails to find any departure from the null model of Brownian evolution at the 5% level of statistical significance (Figure 3c, d). These results are covered in more detail in the discussion.

The final example is taken from (Feilich, 2016) who investigated the evolution of body shape, caudal fin shape, dorsal fin shape, and anal fin shape in 131 African cichlid fishes. Reanalysing the data for anal fin shape using the pointwise envelope (Figure 3e) confirms the spike in relative disparity coinciding with the Cichlinae–Pseudocrenilabrinae split 45-75 MYA reported in the original paper as well as the spike nearer to the present day that coincides with the haplochromine radiation (Feilich, 2016). In contrast, the rank envelope method finds no discernable difference (at the 5% level of significance) from the null model of Brownian evolution at any point in the evolutionary timeline (Figure 3f). Re-analysis of body shape, dorsal fin shape, and caudal fin shape evolution using the rank envelope method does retain support for the peaks in disparity associated with the haplochromine radiation found by (Feilich, 2016) using the pointwise envelope method (Figure S2).

## Discussion

The object of this study has been to highlight the unacceptably high false positive rates of the pointwise envelope test, and offer an alternative solution, the global rank envelope test (Myllymaki et al., 2017) that does not have the same multiple testing issues, but which still allows identification of the time period where the non-Brownian trait evolution may have occurred. Envelope tests using the DTT pointwise envelope method continue to be useful and popular (e.g. Johnson and Omland, 2004, Slater et al., 2010, Dornburg et al., 2011, Blackburn et al., 2013, Ingram, 2015, Arbour and Lopez-Fernandez, 2016, Aristide et al., 2016, Feilich, 2016, Hlusko et al., 2016, Weber et al., 2016) as the time periods over which trait evolution has been non-Brownian can often be linked to specific events that may have triggered the burst of trait evolution (e.g. Slater et al., 2010, Feilich, 2016, Hlusko et al., 2016). Unfortunately, the results presented here (Figures 1–3) suggest reinterpretation of some previous analyses may be required using a global envelope test such as that introduced here rather than a pointwise envelope test.

A number of methods beyond the pointwise envelope test have been used and there is clear variability in their ability to detect departures from the null model (Figure 2). The MDI (Harmon et al., 2003) is a global test as it sums up the difference between the empirical DTT and the average of the simulations of the null model. However, the MDI can be relatively insensitive to early bursts in diversity that lead to larger between-clade disparity (Figure 2); can return a high false positive rate for datasets with less than 40 species (Figure 1); and it is possible for a large difference from the null model in one time period to be cancelled out by smaller deviations in the opposite direction at other time periods. However, the MDI might be better at detecting late bursts in diversity (Figure 2). This is possibly because late bursts are often associated with long time periods of disparity concentrated within clades (eg Figure 2a, e) and there is less chance the deviation from the median of the simulations is cancelled out by negative deviations elsewhere. On the other hand, there is little to choose between the rank envelope test and node height test based upon the results presented here (Figure 1, 2). The main advantage of the rank envelope method is that it provides a visualization of how disparity changes over time and it is easier to see where the burst of trait evolution may have occurred. Given this, and the slightly better performance when species numbers are small and/or early/late bursts are weak, the rank envelope method might be preferable.

Tests using three datasets that are freely available show how inferences and conclusions can change in quite important ways when the rank envelope test is used instead of the pointwise envelope (Figure 3). Macroevolutionary investigations have often used multiple methods (eg node height test, MDI test and DTT plots (Slater et al., 2010); MDI test and DTT plots (Arbour and Lopez-Fernandez, 2016)), to test for departures from Brownian evolution, but given the results presented here there is a risk that a mixture of results will be produced. For example, before removing species considered to be outliers, Slater et al. (2010) found evidence supporting an early burst in cetacean body size using the pointwise envelope test (replicated here in Figure 3), but neither the MDI test nor the node height test could find a statistically significant deviation from the null model of Brownian evolution. These discrepancies are expected given the high false positive rates of the pointwise envelope test (Figure 1), and re-analysis using the global rank envelope test shows agreement with the results of the initial node height test, and MDI test before removal of outliers (Slater et al. (2010); Figure 3).

The methods investigated here all use the same hypothesis testing approach. That is to say we test our data against a suitable null model to see if there are detectable departures from the null model. A different approach is to consider a number of candidate models and ask which model best describes the data (Johnson and Omland, 2004). The advantage of this model selection approach is that multiple models are considered simultaneously, but of course there is no guarantee that the best model, usually determined by some information theoretic criterion, is a ‘good’ descriptor of the data. The model selection approach has been developed for trait evolution by Harmon et al. (2010) who used maximum likelihood methods to fit models that could produce Brownian evolution, increasing or decreasing trait diversification rates, as well as selective peaks where the trait value has a tendency to return to a medial value. Using the likelihood ratio test, they found the Brownian evolution and the selective peak (Ornstein-Uhlenbek) models to be the most frequently selected across 49 clades, implying early bursts in trait evolution are relatively rare. Slater and Pennell (2014) extended this method by employing a posterior predictive approach instead of the likelihood ratio test. The posterior predictive approach proceeds by fitting the parameters to the candidate models using maximum likelihood as in (Harmon et al., 2010), but model selection is based upon sampling the trait evolution from the fitted models and then comparing the fit of each model to the observed trait values. Slater and Pennell (2014) developed this method using either the MDI test, or the node height test and showed both of these posterior predictive methods can have a higher power to detect early bursts in trait evolution compared to the maximum likelihood ratio approach of Harmon et al. (2010). Re-analysing the cetacean dataset with these methods led to the conclusion that an early burst model best described the evolution of whale body size (Slater and Pennell, 2014). This is not surprising given the rank envelope test clearly shows the empirical DTT curve is close to falling below the lower confidence interval (Figure 2d). Ultimately, the user needs to choose between the null model testing and model selection methods, but the rank envelope test developed here could easily be incorporated into the posterior predictive methods of Slater and Pennell (2014), since the ranking of the observed DTT curve in the ensemble of simulations from each of the candidate models generates a single metric, the global rank amongst the set of model curves, that could then be used to compare the models.

In summary, the pointwise envelope test method that has been employed to investigate bursts in trait evolution shows unacceptably high type 1 statistical errors and should not be used. The rank envelope test that was introduced by Myllymaki et al. (2017) for spatial point pattern analysis, ranks the extremness of the empirical DTT curve against the suite of realisations of an appropriate null model, and is instead shown to possess the desirable statistical properties. This method is shown to be at least as powerful as other popular methods for detecting bursts of trait evolution, whilst retaining the advantages of graphical interpretation of where in time the significant bursts of evolution occur. Null model testing is a commonly used tool in ecology and evolution (eg Gotelli and Ulrich (2012)) and the rank envelope test is a flexible method that allows inference without the problem of multiple testing when the null model returns a functional relationship (ie a vector) rather than a singular value (ie a scalar quantity).

## Acknowledgements

This work was supported by the Engineering and Physical Sciences Research Council grant EP/N007336/1. The author would like to thank Alex Pigot who provided useful suggestion and comments on earlier versions of the manuscript.

---

**Figure S1.**
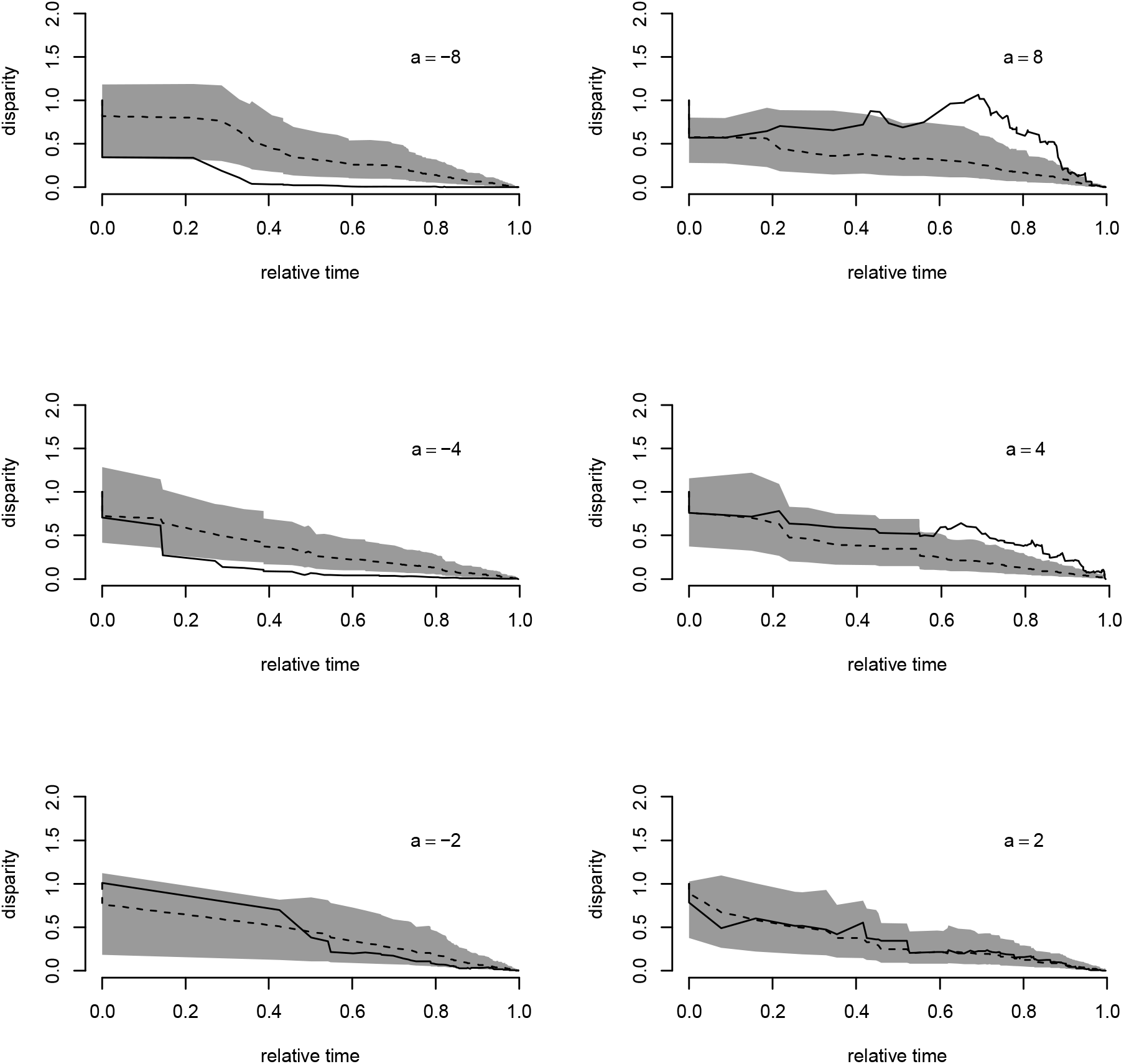
Examples of simulated data sets for disparity through time analyses where the rate of trait evolution either slows down (*a* < 0) or speeds up (*a* > 0) over time (black solid lines). Broken lines represent the median disparity through time for 2500 simulations of the null model of Brownian evolution (ie *a* = 0), and the shaded region is the 95% confidence interval for the simulations based upon the rank envelope method (see main text). Simulations are for simulated phylogenetic trees with 100 species at the tips.

**Figure S2.**
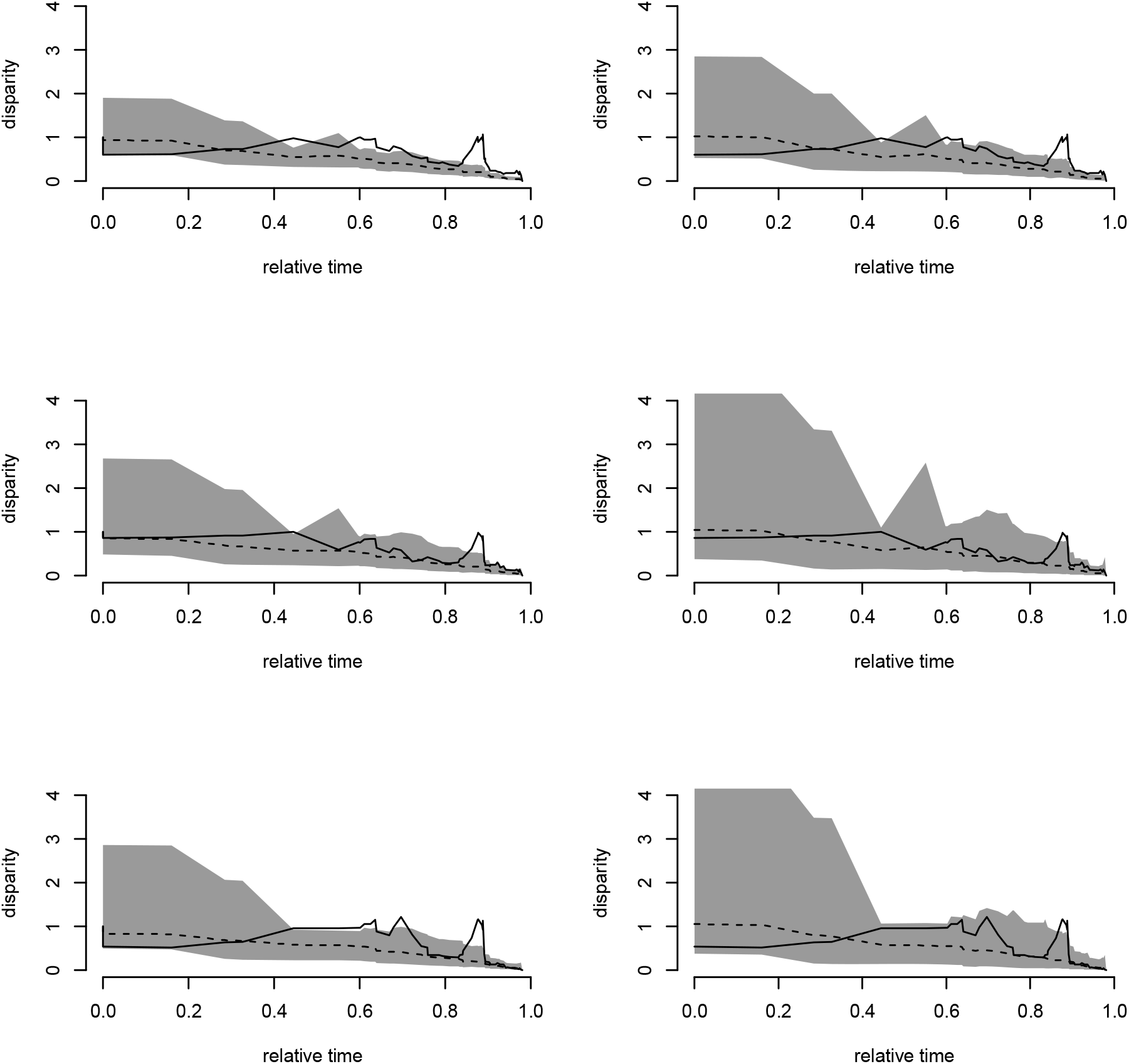
Comparisons of inference from using the (left hand column) pointwise envelope test and the (right hand column) rank envelope test for trait disparity through time for (top row) body shape; (middle row) caudal fin shape; and (bottom row) dorsal fin shape, for 131 species of African cichlid fishes. Data is taken from (Feilich, 2016). In each panel the empirical pattern (solid black line) is compared to the median of 5000 simulations of the null model of Brownian evolution of the trait values (broken lines) on the phylogenetic tree, and the shaded regions correspond to the 95% confidence region.

